# CRISPR Screen in Regulatory T Cells Reveals Ubiquitination Modulators of Foxp3

**DOI:** 10.1101/2020.02.26.966911

**Authors:** Jessica T. Cortez, Elena Montauti, Eric Shifrut, Yusi Zhang, Oren Shaked, Yuanming Xu, Theodore L. Roth, Dimitre R. Simeonov, Yana Zhang, Siqi Chen, Zhongmei Li, Ian A. Vogel, Grace Y. Prator, Bin Zhang, Youjin Lee, Zhaolin Sun, Igal Ifergan, Frédéric Van Gool, Jeffrey A. Bluestone, Alexander Marson, Deyu Fang

**Affiliations:** Biomedical Sciences Graduate Program, University of California, San Francisco, California 94143, USA; Department of Microbiology and Immunology, University of California, San Francisco, California 94143, USA; Diabetes Center, University of California, San Francisco, California 94143, USA; Innovative Genomics Institute, University of California, Berkeley, California 94720, USA; Department of Pathology; Northwestern University Feinberg School of Medicine, 303 E. Chicago Ave, Chicago, IL 60611; Department of Immunology; the Fourth Military Medical University, Xi’an 710032, China; Robert H. Lurie Comprehensive Cancer Center, Department of Medicine-Division of Hematology/Oncology, Northwestern University Feinberg School of Medicine, Chicago, IL 60611, USA; Department of Pharmacology, Dalian Medical University School of Pharmacy, Dalian 116044, China; Department of Microbiology and Immunology; Northwestern University Feinberg School of Medicine, 303 E. Chicago Ave, Chicago, IL 60611, USA; UCSF Helen Diller Family Comprehensive Cancer Center, University of California, San Francisco, California 94158, USA; Department of Medicine, University of California, San Francisco, California 94143, USA; Chan Zuckerberg Biohub, San Francisco, California 94158, USA; Rosalind Russell/Ephraim P. Engleman Rheumatology Research Center, University of California, San Francisco, San Francisco, CA, 94143, USA

## Abstract

Regulatory T cells (Tregs) are required to control immune responses and maintain homeostasis but are a significant barrier to anti-tumor immunity^1^. Conversely, Treg instability, characterized by loss of the master transcription factor Foxp3 and acquisition of pro-inflammatory properties^2^, can promote autoimmunity and/or facilitate more effective tumor immunity^3,4^. A comprehensive understanding of the pathways that regulate Foxp3 could lead to more effective Treg therapies for autoimmune disease and cancer. Despite improved functional genetic tools that now allow for systematic interrogation, dissection of the gene regulatory programs that modulate Foxp3 expression has not yet been reported. In this study, we developed a CRISPR-based pooled screening platform for phenotypes in primary mouse Tregs and applied this technology to perform a targeted loss-of-function screen of ∼490 nuclear factors to identify gene regulatory programs that promote or disrupt Foxp3 expression. We discovered several novel modulators including ubiquitin-specific peptidase 22 (Usp22), Ataxin 7 like 3 (Atxn7l3) and ring finger protein 20 (Rnf20). Members of the deubiquitination module of the SAGA chromatin modifying complex, Usp22 and Atxn7l3, were discovered to be positive regulators that stabilized Foxp3 expression; whereas the screen suggested Rnf20, an E3 ubiquitin ligase, is a negative regulator of Foxp3. Treg-specific ablation of Usp22 in mice reduced Foxp3 protein and created defects in their suppressive function that led to spontaneous autoimmunity but protected against tumor growth in multiple cancer models. Foxp3 destabilization in Usp22-deficient Tregs could be rescued by ablation of Rnf20, revealing a reciprocal ubiquitin switch in Tregs. These results reveal novel modulators of Foxp3 and demonstrate a screening method that can be broadly applied to discover new targets for Treg immunotherapies for cancer and autoimmune disease.

## Main

While unstable Foxp3 expression in Tregs can result in autoimmunity, similar changes that reduce Treg suppressive function can contribute to more effective anti-tumor immune responses^4^. Understanding the fundamental regulators of Foxp3 is critical, especially as we navigate new potential applications for Treg therapies to treat autoimmunity and cancer^5^.

To discover novel regulators of Foxp3 stability, we developed a pooled CRISPR screening platform in primary mouse Tregs (**Fig. 1a**). We first designed a targeted library of ∼490 nuclear factors based on optimized single guide RNA (sgRNA) sequences from the Brie library^6^ (**Extended Data Fig. 1a**) and used a retroviral vector to introduce this library into *ex vivo* Tregs isolated from *Foxp3*^GFP-Cre^*Rosa26*^LSL-RFP^*Cas9* mice (**Extended Data Figs. 1b-1e**). We then stained for endogenous Foxp3 protein and sorted the highest Foxp3-expressing cells (Foxp3^high^) and the lowest (Foxp3^low^). MAGeCK software^7^ systematically identified sgRNAs that were enriched or depleted in Foxp3^low^ cells relative to Foxp3^high^ cells (**<u>Supplementary Table 1</u>**). We were able to maintain high sgRNA coverage of our library (∼1000x) and non-targeting (NT) control sgRNAs showed no effect (**Extended Data Fig. 1f, 1g**) which provided confidence that our hits identified biological pathways controlling Foxp3 levels.

**Figure 1.**
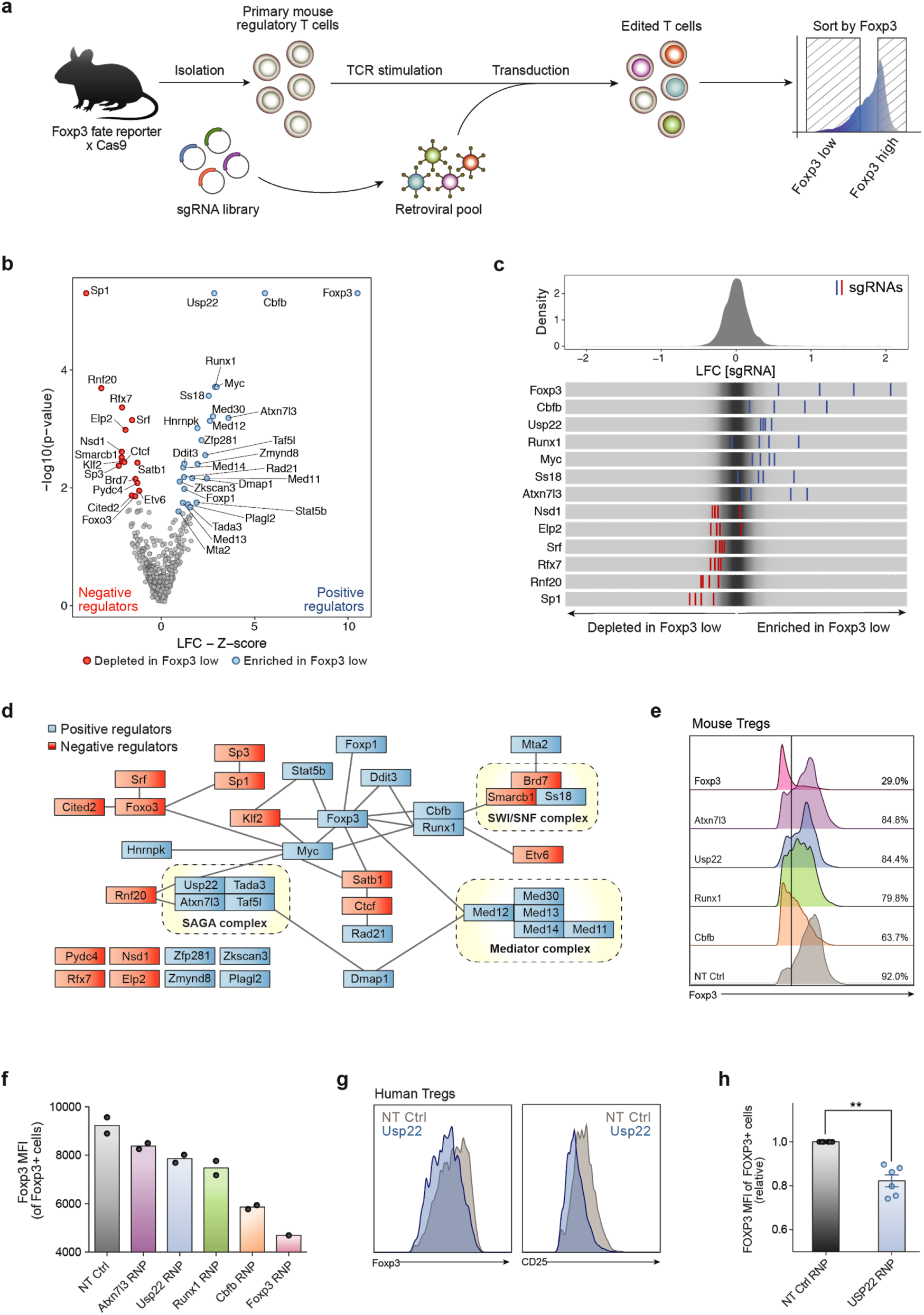
Discovery and Validation of Regulators of Foxp3 in Primary Mouse Tregs Using a Targeted Pooled CRISPR Screen. a) Diagram of pooled CRISPR screening platform in primary mouse Treg cells. b) Volcano plot for hits from the screen. X-axis shows Z-score for gene-level log2 fold-change (LFC); median of LFC for all single guide RNAs (sgRNAs) per gene, scaled. Y-axis shows the p-value as calculated by MAGeCK. Red are negative regulators (depleted in Foxp3 low cells), while blue dots show all positive regulators (enriched in Foxp3 low cells) defined by FDR < 0.5 and Z-score > 0.5. c) Top panel: distribution of sgRNA-level LFC values of Foxp3 low over Foxp3 high cells for 2,000 guides. Bottom panel: LFC for all four individual sgRNAs targeting genes enriched in Foxp3 low cells (blue lines) and depleted genes (red lines), overlaid on grey gradient depicting the overall distribution. d) Schematic of experimentally determined and predicted protein-protein interactions between top hits, 16 negative regulators (red) and 25 positive regulators (red), generated by STRING-db. Black lines connect proteins that interact and dotted lines depict known protein complexes. e) Foxp3 expression 5 days post electroporation of Cas9 RNPs in mouse Tregs as measured by flow cytometry of top screen hits. Histograms depict cells that have been pre-gated on Foxp3+ cells. f) Mean fluorescence intensity (MFI) of Foxp3 from data in panel e. g) Representative histogram showing MFI of FOXP3 and CD25 from human Tregs. h) Statistical analysis of FOXP3 MFI from human Tregs in 6 biological replicates. **P < 0.01.

Our screen revealed many novel Foxp3 regulators, with a bias towards identifying positive regulators over negative regulators (**Figs. 1b, 1c**). sgRNAs enriched in the Foxp3^low^ population reflect positive regulators (blue) that promote Foxp3 expression while sgRNAs depleted in the Foxp3^low^ population reflect negative regulators that inhibit Foxp3 expression (red). We identified many established regulators known to be important for maintenance of Foxp3 expression including Foxp3 itself, Cbfb, Runx1 and Stat5b^8–14^ as positive regulators and Sp3 and Satb1 as negative regulators^9,15^ providing further confidence in our hits. Importantly, several novel factors and complexes that modulate Foxp3 were identified including positive regulators Usp22, Atxn7l3 and negative regulator Rnf20. The deubiquitinase (DUB) Usp22 and cofactor Atxn7l3, are both members of deubiquitination module of the SAGA chromatin modifying complex^16^ (**Fig. 1d**).

To validate the phenotype of our screen hits, we assessed five of the top-ranking positive regulators by individual CRISPR knockout with Cas9 ribonucleoproteins^17^ (RNPs) (**Extended Data Fig. 2a**). All guides tested resulted in a decrease in Foxp3 expression reproducing the screen data (**Figs. 1e, 1f and Extended Data Fig. 2e**). These results confirmed the candidate genes identified from our screen as positive regulators of Foxp3. As our screen indicated that Usp22 is a positive regulator of Foxp3, we next wanted to assess the potential therapeutic relevance of USP22 by knocking it out with RNPs in human Tregs. We saw a significant decrease in FOXP3 and CD25 mean fluorescence intensity (MFI) (**Figs. 1g,h)** and frequencies of FOXP3^hi^CD25^hi^ cells in USP22-deficient human Tregs (**Extended Data Figs. 2b-d**) across six biological replicates.

To understand the *in vivo* significance of Usp22 in Tregs, we generated mice with Treg-specific ablation of Usp22 by creating *Usp22*^*fl/fl*^ mice (**Extended Data Fig. 3a, 3b**) and crossing them with *Foxp3*^*YFP-Cre*^ mice^18^. Western blot analysis confirmed specific deletion of Usp22 in Treg cells, but not in CD4^+^ conventional T (Tconv) cells (**Extended Data Fig. 3c**). *Usp22*^*fl/fl*^*Foxp3*^*YFP-Cre*^ knockout (KO) mice had a marked decrease in Foxp3 MFI and frequency in Tregs isolated from spleens, thymus and peripheral lymph nodes (pLN) compared to littermate *Usp22*^*+/+*^*Foxp3*^*YFP-Cre*^ (WT) mice (**Figs. 2a-c, Extended Data Fig. 3d**). Western blot analysis confirmed a significant reduction in Foxp3 protein in Usp22-null Tregs **(Fig. 2d)**. A decrease in Foxp3^+^ cells was also seen in induced Tregs (iTregs), although less pronounced with increasing levels of TGF-β (**Extended Data Fig. 3e**). Given the diminished Foxp3 levels in Usp22 KO Tregs, we reasoned that these cells may exhibit defects in suppressive function. Indeed, Usp22 KO Tregs were less able to suppress T effector cells than WT Tregs (**Fig. 2e, 2f**) suggesting that Usp22 is critical to maintain Treg suppressive function. These data substantiate our screen data and suggest Usp22 is critical to maintain Treg stability by promoting Foxp3 levels.

**Figure 2.**
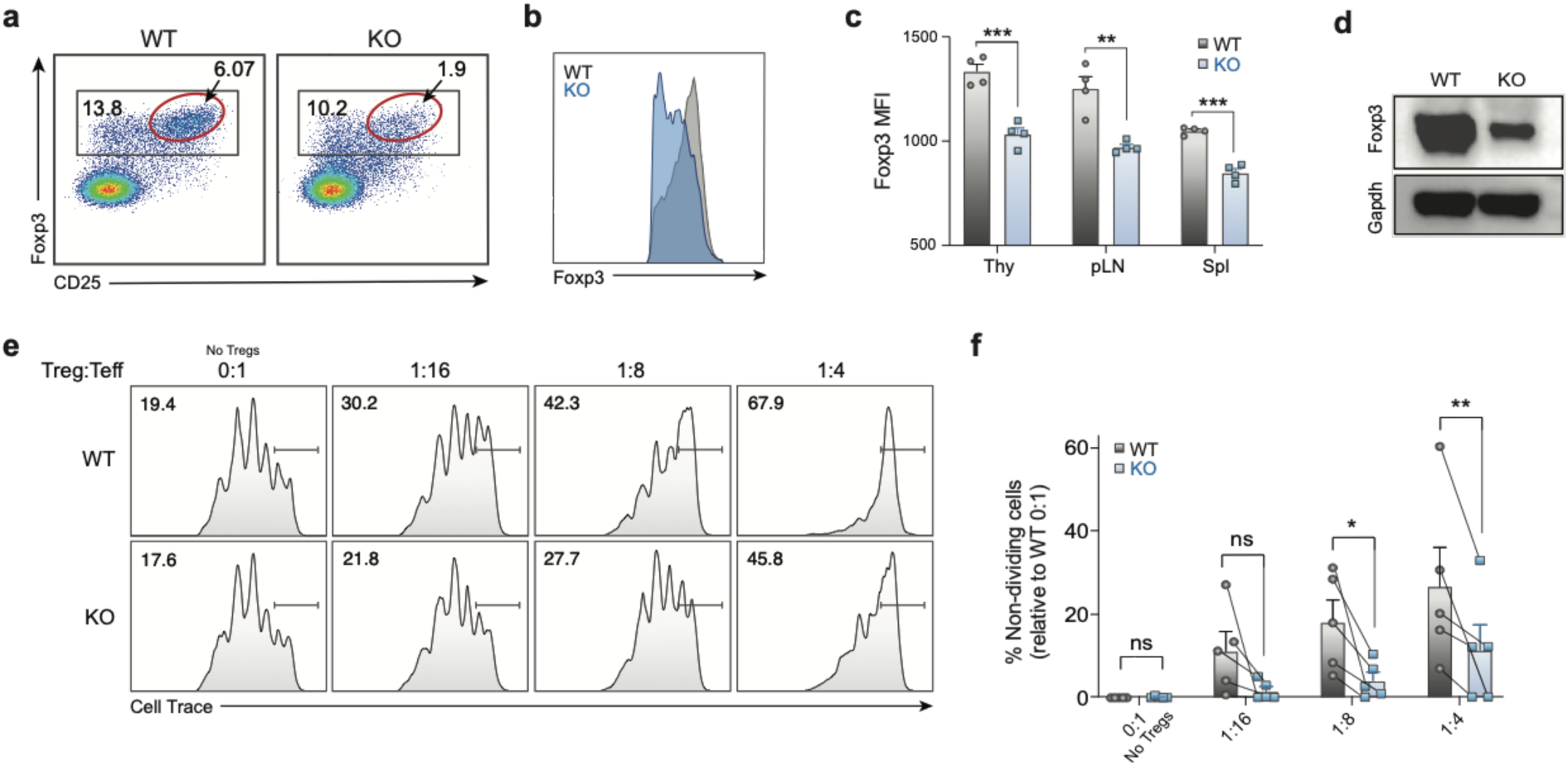
Usp22 is required to maintain Treg suppressive function and Foxp3 stability *in vivo.* a) Representative flow cytometry analysis of the Treg population (gated on CD4^+^ cells) from the spleen of Treg-specific Usp22 WT and KO mice. The Foxp3^hi^ CD25^hi^ subpopulation is highlighted with a red gate. b) Histogram of Foxp3 expression from the spleen of Treg-specific Usp22 WT and KO mice from panel a. c) Statistical analysis of Foxp3 MFI from thymus (Thy), peripheral lymph nodes (pLN) and spleen (Spl). *P < 0.05, **P < 0.01, ***P < 0.001, ****P < 0.0001. d) Western blot analysis of Foxp3 protein level from Tregs isolated from spleen and LN in Usp22 WT and KO mice. Gapdh was used as a loading control. e) *In vitro* suppressive activity of Tregs assessed by the division of naïve CD4^+^CD25^-^ T cells. Naïve cells were labeled with cytosolic cell proliferation dye and activated by anti-CD3 and antigen presenting cells (irradiated splenocytes from wild-type mice, depleted of CD3+ T cells), then co-cultured at various ratios (as indicated above) with YFP+ Treg cells sorted from 8-week-old Usp22 WT or KO mice. Numbers indicate the percentage of non-dividing cells for each ratio. f) Summary data of *in vitro* suppression experiments. *P < 0.05, **P < 0.01, ***P < 0.001, ****P < 0.0001.

Control of Foxp3 stability, and thus Treg function, can occur at both the transcriptional and post-translational level^19^. Epigenetic modifications at the *Foxp3* locus and at Treg-associated loci can affect Foxp3 transcription^20–25^. Foxp3 protein can also be dynamically controlled post-translationally by DUBs or ubiquitin ligases in response to proinflammatory signals^26–30^. As Usp22 is a component of the chromatin modifying SAGA complex, we hypothesized that Usp22 controls Foxp3 expression through transcriptional regulation. IRES-YFP knock-in to the *Foxp3* locus of these mice allowed us to use YFP as a reporter to assess the effect of Usp22 on *Foxp3* transcript levels. Similar to endogenous Foxp3 protein, YFP MFI was significantly decreased in Usp22-null Tregs isolated from the thymus, pLN and spleen in KO mice compared to littermate WT mice **(Figs. 3a-c)**, despite normal Treg frequencies (**Extended Data Fig. 5a**). Furthermore, by qPCR Foxp3 mRNA transcripts were significantly reduced in KO spleens compared to littermate WT mice **(Fig. 3d)**. RNA sequencing also confirmed that *Foxp3* transcripts are significantly reduced in Usp22 KO cells relative to WT **(Fig. 3e)**. These results are consistent with Usp22 tuning Foxp3 expression at the transcriptional level.

**Figure 3.**
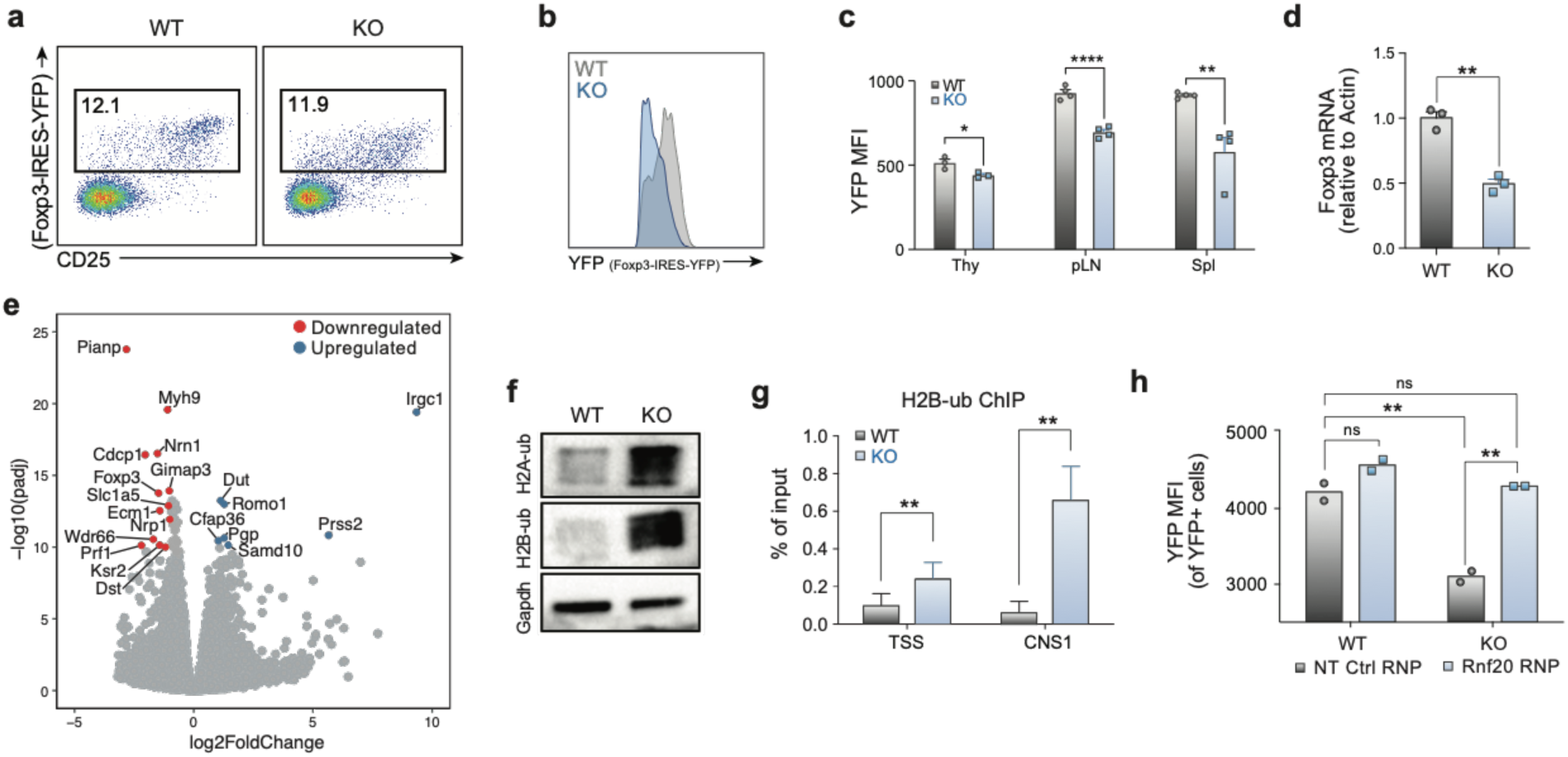
Usp22 Regulates Foxp3 Transcription by Reciprocal Ubiquitin Modulators. a) Representative flow cytometry analysis of the YFP^+^ Treg population (gated on CD4^+^ cells) from the spleen and lymph nodes of Treg-specific Usp22 WT and KO mice. b) Histogram of YFP expression from the spleen and lymph nodes of Treg-specific Usp22 WT and KO mice from panel a. c) Statistical analysis of YFP MFI from thymus (Thy), peripheral lymph nodes (pLN) and spleen (Spl), n=3-4. d) qPCR analysis mRNA level of Foxp3 in sorted YFP+ cells of spleen from Usp22 WT and KO mice. ns indicates no significant difference, *P < 0.05, **P < 0.01, ***P < 0.001 by two-tailed unpaired t-test. e) Volcano plot for RNA sequencing of Usp22 WT and KO Tregs. X-axis shows log2FoldChange (LFC). Y-axis shows the –log10 of the adjusted p-value (padj) as calculated by DESeq2. Genes downregulated in the KO are shown in red and genes upregulated are shown in blue defined by padj <1e-10 and LFC > 1. f) Western blot analysis of ubiquitinated histone 2A (H2A-ub) and ubiquitinated histone 2B (H2B-ub) from iTregs from Usp22 WT and KO mice. Gapdh was used as a loading control. g) ChIP-qPCR data analysis for H2B-ub where primers amplified across the transcriptional start site (TSS) and the CNS1 enhancer region of the *Foxp3* locus. Data are normalized to the input. h) Representative analysis of reciprocal regulation of Foxp3 by deubiquitinase Usp22 and E3 ubiquitin ligase Rnf20. Statistical analysis of Foxp3 MFI in Usp22 WT and KO Tregs electroporated with either NT control or Rnf20 RNP. ns indicates no significant difference, **P < 0.01 by two-way ANOVA followed by Holm–Sidak multiple comparisons test.

Usp22 is required for SAGA-mediated deubiquitination of histones, which regulates transcriptional activity^31^. We therefore tested if histone ubiquitination was altered in Usp22 KO Tregs. Western blot analysis confirmed that KO mice had increased levels of ubiquinated histone 2A and histone 2B (H2A-ub and H2B-ub, respectively) in iTregs compared to WT (**Fig. 3f**). ChIP-qPCR confirmed increased H2B-ub at the *Foxp3* transcriptional start site (TSS) and the conserved non-coding sequence 1 (CNS1) enhancer^22^ in the *Foxp3* locus in KO iTregs (**Fig. 3g and Extended Data Fig. 5b**). We similarly saw increased H2A-ub at the *Foxp3* locus, but this result was less pronounced than H2B-ub (**Extended Data Fig. 5c**). Our screen also nominated E3-ubiquitin ligase Rnf20 as a negative regulator of Foxp3. We hypothesized that the DUB Usp22 and E3-ubiquitin ligase Rnf20 might have an epistatic relationship given their reciprocal effects on histone ubiquitination. To test this, we used RNPs to knockout Rnf20 in Usp22 KO or WT Tregs. In Usp22-deficient Tregs, Rnf20 RNP knockout was able to rescue the impairment in *Foxp3* transcript levels **(Fig. 3h)**. Taken together, these results revealed reciprocal regulation of *Foxp3* transcripts by Usp22 and Rnf20.

Given the reported function of Usp22 as a DUB^20^, we investigated whether Foxp3 protein can be directly targeted by Usp22. In addition to the observed effects on *Foxp3* transcript levels, Usp22 loss contributed to increased Foxp3 ubiquitination and degradation (**Extended Data Fig. 4**). These results suggest that Usp22 can regulate both Foxp3 transcript and protein.

To determine the *in vivo* functional relevance of Usp22 deficiency in Tregs, we characterized the spontaneous autoimmune symptoms of Usp22 KO mice. While *Usp22*^*fl/fl*^*Foxp3*^*YFP-Cre*^ mice were born at normal size, their body weight became significantly reduced compared to age and sex matched WT control mice after 5 weeks of age (**Fig. 4a**). This reduction in body weight was not due to YFP-Cre knock-in expression, since the body weight of 8-week-old C57BL/6 WT mice with or without YFP-Cre are comparable, and body weight reduction was only observed in *Usp22*^*fl/fl*^*Foxp3*^*YFP-Cre*^ KO mice (**Extended Data Fig. 6a**). We next assessed whether this body weight reduction might be due to chronic inflammation as is observed with impaired Treg function^1^. Indeed, flow cytometry analysis detected greater frequencies and increased absolute numbers of effector memory T cells (CD44^hi^CD62L^lo^) and corresponding lower percentages and absolute numbers of naïve T cells (CD44^lo^CD62L^hi^) in 7-month-old KO mice compared to littermate WT control mice **(Extended Data Figs. 6b, 6c)**. Additionally, histological analysis of aged mice detected lymphocyte infiltration in multiple organs, including kidney, lung, colon and liver **(Fig. 4b)**. We then assessed Treg suppressive activity *in vivo* using an adoptive transfer model of colitis and a MOG-induced experimental autoimmune encephalomyelitis (EAE) model. In the colitis model, mice that received defective Usp22 KO Tregs were not protected against colitis, in contrast to those that received WT Tregs **(Figs. 4c, 4d)**. Similarly, in the EAE model, Usp22 KO mice showed worse clinical scores compared to WT mice suggesting an inability of the Usp22-deficient Tregs to prevent autoimmunity **(Fig. 4e)**.

**Figure 4.**
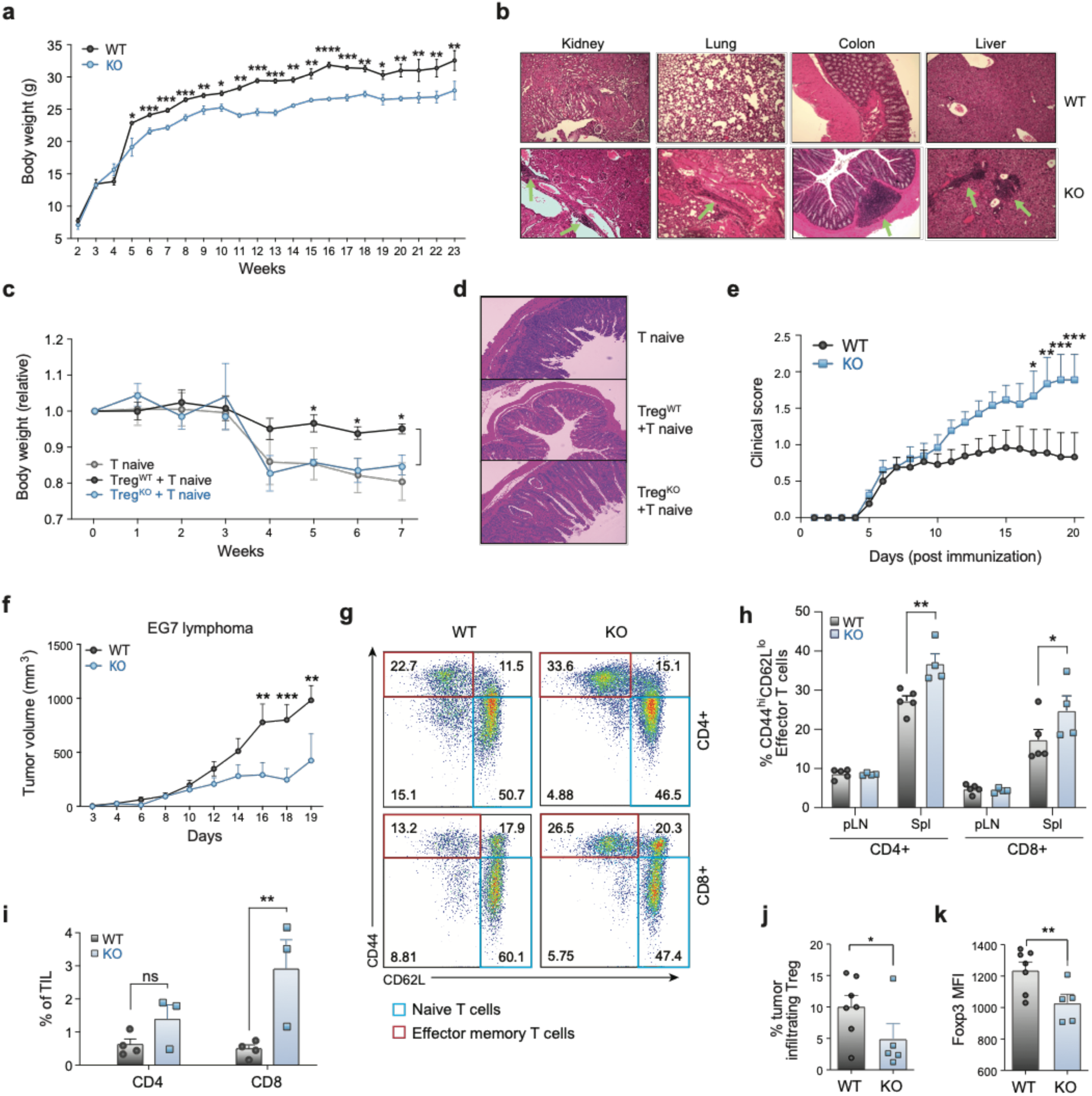
Treg-Specific Ablation of Usp22 Results in Spontaneous Autoimmune Symptoms and Enhances Anti-Tumor Immunity. a) Body weight differences between Usp22 WT and KO littermate mice. b) Hematoxylin-and-eosin (H&E) staining of kidney, lung, colon and liver sections from 7-month-old WT and KO mice. Original magnification, x100. c) Weight of Rag^-/-^ recipient mice (n=4-5 per group) over time after adoptive transfer of CD4^+^CD25^-^CD44^lo^CD62^hi^ (CD45.1^+^) naïve T cells sorted from SJL mice alone or together with CD4^+^YFP^+^ (CD45.2^+^) Treg cells from 9-week-old Usp22 WT or KO mice, presented relative to weight at day 0, set as 1. d) H&E staining of colon tissues from the Rag^-/-^ recipient mice shown in panel c, 7 weeks post transfer. Original magnification at 100x (fold). e) Clinical severity of EAE in Usp22 WT and KO mice (n=4) was monitored until day 20 post immunization with MOG peptide. f) EG7 lymphoma tumor growth is inhibited in Usp22 KO mice. Mice (n=4 or 5 per group) were subcutaneously inoculated with 106 EG7 cells. The tumor volume was measured every 2 or 3 days by scaling along 3 orthogonal axes (x, y, and z) and calculated as (xyz)/2. g) Representative flow cytometry analysis of the expression of CD44 and CD62L in both CD4+ and CD8+ T cells of spleen from EG7 tumor-bearing mice. h) The percentage of effector memory-like activated T cells (CD44^hi^CD62L^lo^) from EG7 tumor-bearing mice summarized. i) Statistical analysis of tumor-infiltrating lymphocyte (TIL) percentages from EG7-bearing Usp22 WT or KO mice after tumor inoculation. j) Statistical analysis of tumor-infiltrating Treg percentages from EG7-bearing Usp22 WT or KO mice after tumor inoculation. k) Foxp3 MFI of the CD4+Foxp3+ EG7 tumor-infiltrating Treg population summarized, (n=5-7). Data are mean ±SEM and are representative of three independent experiments. ns indicates no significant difference, *P < 0.05, **P < 0.01, ***P < 0.001, ****P < 0.0001.

Since these data suggest that Usp22 deficiency reduces Foxp3 stability and impairs Treg suppressive function, we next tested whether Usp22 KO mice would exhibit increased anti-tumor immunity using syngeneic tumor models. As expected, EG7 lymphoma tumors were significantly inhibited by Treg-specific Usp22 gene deletion **(Fig.4f)**. We next examined the immune responses in these tumor-bearing mice and found greater proportions of both effector-memory CD4^+^ and CD8^+^ T cells in the spleens of KO mice compared to WT mice **(Figs. 4g, 4h)**. Further analysis of tumor-infiltrating lymphocytes indicated a significant increase in CD8^+^ T cells in EG7 tumor-bearing KO mice **(Fig. 4i)** and subsequent decrease in intratumoral Tregs **(Fig. 4j)**. Consistent with the lymphoid organs, we found that the Foxp3 MFI was significantly decreased in the intratumoral Treg cells from EG7 tumor-bearing KO mice **(Fig. 4k)**. Taken together, these data indicate Usp22 KO impairs Treg suppressive function and reduces Treg abundance in tumors, consequently enhancing the anti-tumor immune response. Similarly, we show that Usp22 KO mice exhibit increased anti-tumor immunity in multiple tumor models **(Extended Data Figs. 7a-c)**. These results highlight Usp22 in Tregs as a new potential target for anti-tumor immunotherapies.

Here, we developed the first CRISPR-based pooled screening platform for primary mouse Tregs and applied this technology for systematic identification of gene modifications that control Foxp3 levels. We discovered several novel regulators of Foxp3 including Usp22, Atxn7l3 and Rnf20. We developed a Treg-specific Usp22 KO mouse and showed that Usp22 is critical to stabilize Foxp3 and maintain suppressive functions *in vivo*. We demonstrate that Usp22 is a regulator of *Foxp3* transcript levels, likely through deubiquitination of H2B at the *Foxp3* locus, and that Usp22 can also regulate Foxp3 post-translationally. Mice with Usp22-null Tregs showed an enhanced anti-tumor immune response and inability to resolve autoimmune inflammation. This study provides a resource of novel Foxp3 regulators that can be perturbed to fine tune Treg function and specifically defines the function of Usp22 and Rnf20 as important modulators of Foxp3 and potential targets for Treg immunotherapies.

## Supporting information

STable1_ScreenData

STable2_Oligos

STable3_Antibodies

STable4_RNASeqData

---

**Extended Data Fig. 1.**
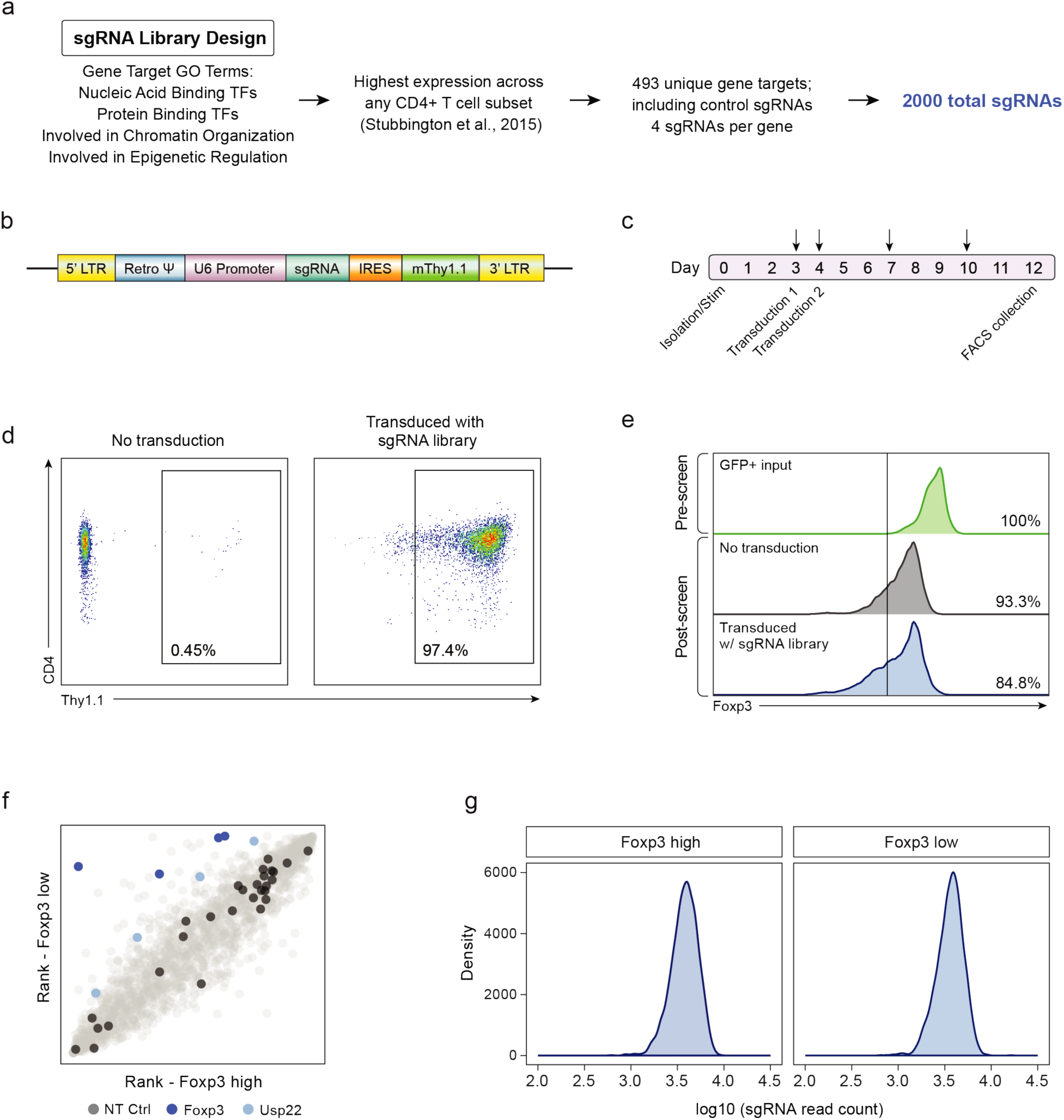
Design and Quality Control of Targeted Pooled CRISPR Screen in Primary Mouse Tregs. a) Design strategy for selection of genes for unbiased targeted library of 493 targets, including 490 nuclear factors and 3 control targets (NT, GFP, and RFP). Genes were selected based on gene ontology (GO) annotation and then sub-selected based on highest expression across any CD4 T cell subset for a total of 2,000 sgRNAs. b) Diagram of MSCV expression vector with Thy1.1 reporter used for retroviral transduction of the sgRNA library. c) Detailed timeline schematic of the 12-day targeted screen pipeline. Arrows indicate when the cells were split and media was replenished. d) Retroviral transduction efficiency of the targeted library in primary mouse Tregs shown by Thy1.1 surface expression measured by flow cytometry. The infection was scaled to achieve a high efficiency multiplicity of infection. e) Foxp3 expression from screen input, output and control cells measured by flow cytometry. Top: Foxp3 expression from input Foxp3^+^ purified Tregs as measured by GFP expression on Day 0. Middle: Foxp3 expression as measured by endogenous intracellular staining from control Tregs (not transduced with library) on Day 12. Bottom: Foxp3 expression as measured by endogenous intracellular staining from screen Tregs (transduced with library) on Day 12. f) Targeted screen (2,000 guides) shows that sgRNAs targeting Foxp3 and Usp22 were enriched in Foxp3 low cells (blue). Non-targeting control (NT Ctrl) sgRNAs were evenly distributed across the cell populations (black). g) Distribution of read counts after next generation sequencing of sgRNAs of sorted cell populations, Foxp3^high^ and Foxp3^low^.

**Extended Data Figure 2.**
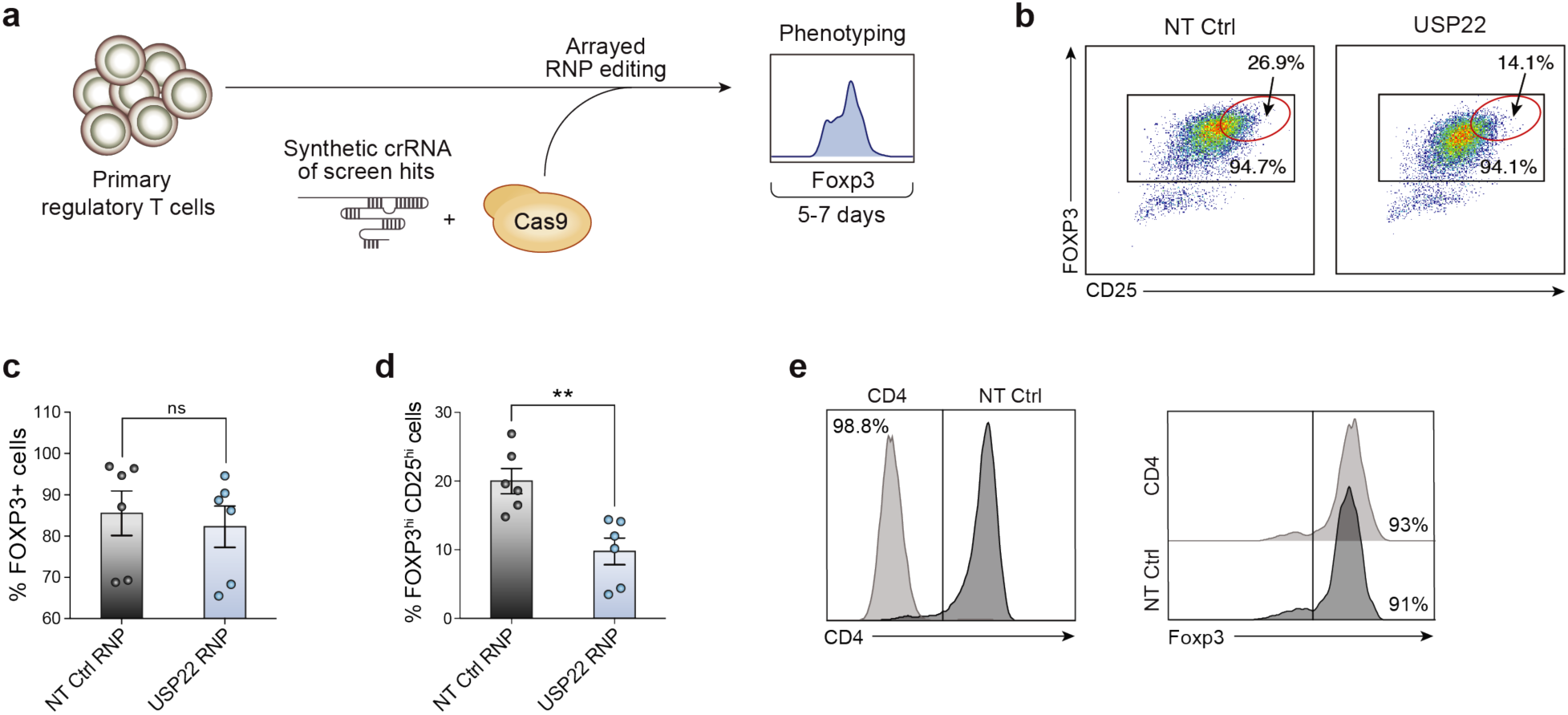
Validation of Gene Targets That Regulate Foxp3 Expression in Primary Mouse and Human Tregs Using RNP Arrays. a) Overview of orthogonal validation strategy using arrayed electroporation of Cas9 RNPs in Tregs. b) Representative flow plots depicting FOXP3 and CD25 expression 7 days post electroporation of Cas9 RNPs in human Tregs. The Foxp3^hi^ CD25^hi^ subpopulation is highlighted with a red gate. c) Percentage of FOXP3^+^ cells from human Tregs in 6 biological replicates. d) Percentage of FOXP3^hi^CD25^hi^ cells from human Tregs in 6 biological replicates. e) RNP controls in mouse Tregs collected 5 days post electroporation. Left: CD4 expression from CD4 RNP (cutting control) compared to NT control. Right: Foxp3 expression from CD4 knockout cells (left panel) compared to NT control.

**Extended Data Fig. 3.**
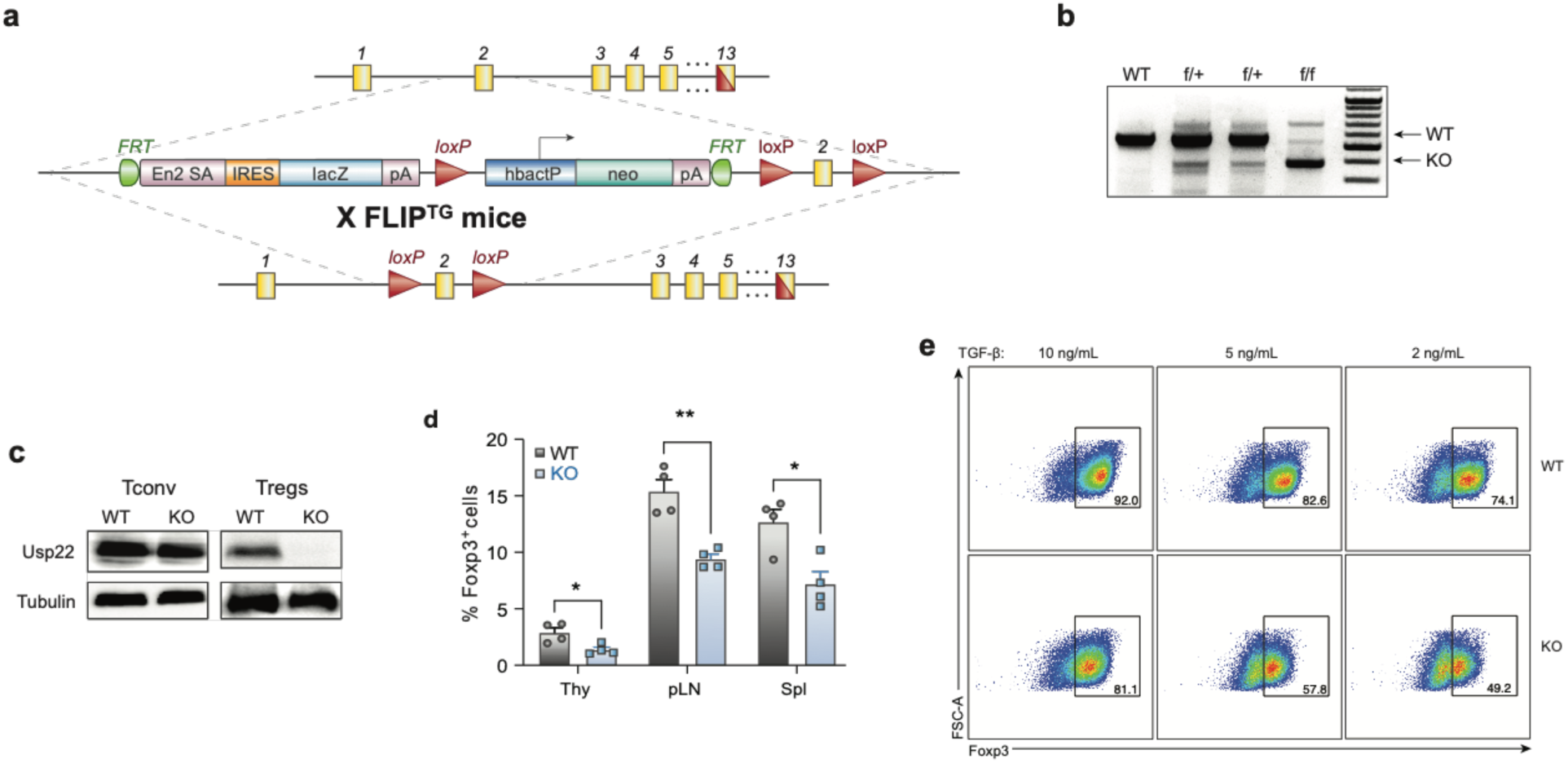
Design and Validation of Treg-specific Usp22 Knockout Mice. a) Diagram of the murine *Usp22* locus. Targeting vector contains IRES-lacZ and a neo cassette inserted into exon 2. b) Genotyping by PCR showed a 600-bp band for the wild-type allele and a 400-bp band for mutant allele, simultaneously in the homozygous floxed (f/f) mice. c) Western blot analysis of Usp22 in CD4^+^CD25^-^ conventional T cells (Tconv) and CD4^+^CD25^+^ Treg cells isolated from WT and KO mice. Tubulin was used as a loading control. d) Statistical analysis of CD4^+^CD25^+^Foxp3^+^ Treg frequencies, n=4, corresponding to Figure 2c. e) iTreg differentiations from WT and KO naïve CD4^+^ T cells with titration of TGF-β.

**Extended Data Fig. 4.**
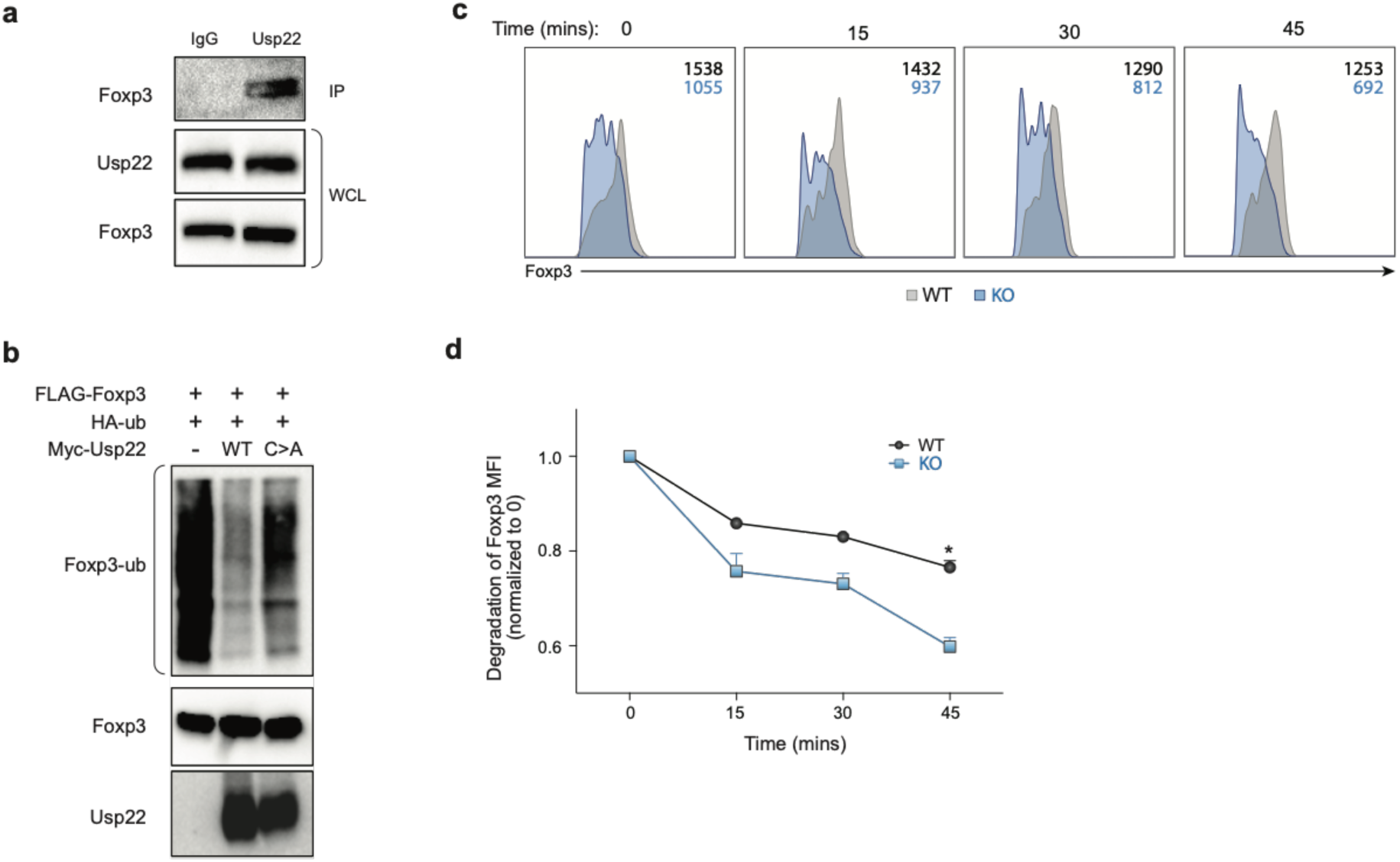
Usp22 Acts as Deubiquitinase to Control Post-Translational Foxp3 Expression. a) Endogenous interaction of Usp22 and Foxp3 in murine iTreg cells from WT mice. Rabbit anti-Usp22 antibody was used to perform the immunoprecipitation and mouse anti-Foxp3 antibody was used to detect the bound Foxp3. Normal rabbit IgG was used as control. b) Ubiquitination assay of Foxp3 with WT or mutant Usp22, where the conserved cysteine (C) residue in the C19 peptidase domain was replaced by an alanine (A) residue. c) Splenocytes isolated from Usp22 WT or KO mice were treated with 200 μg/ml cycloheximide (CHX) for the indicated time course. Inset numbers for each histogram indicate the MFI of Foxp3 (black=WT, blue=KO). d) Foxp3 MFI from the CD4^+^CD25^+^Foxp3^+^ Treg population (n=3). Data are mean ±SEM and are representative of three independent experiments. *P < 0.05.

**Extended Data Fig. 5.**
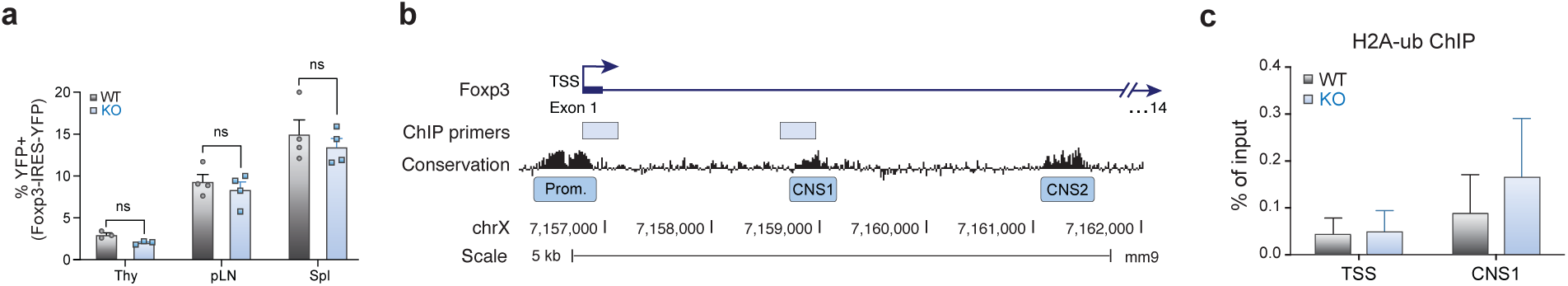
Usp22 Regulates Foxp3 through Transcriptional Mechanisms. a) Statistical analysis of CD4^+^YFP^+^ Treg frequencies, corresponding to Figure 3c. b) Schematic of *Foxp3* locus depicting PCR products used for ChIP-qPCR. c) ChIP-qPCR data analysis for H2A-ub for PCR across the transcriptional start site (TSS) and across the CNS1 enhancer region. Data are normalized to the input.

**Extended Data Fig. 6.**
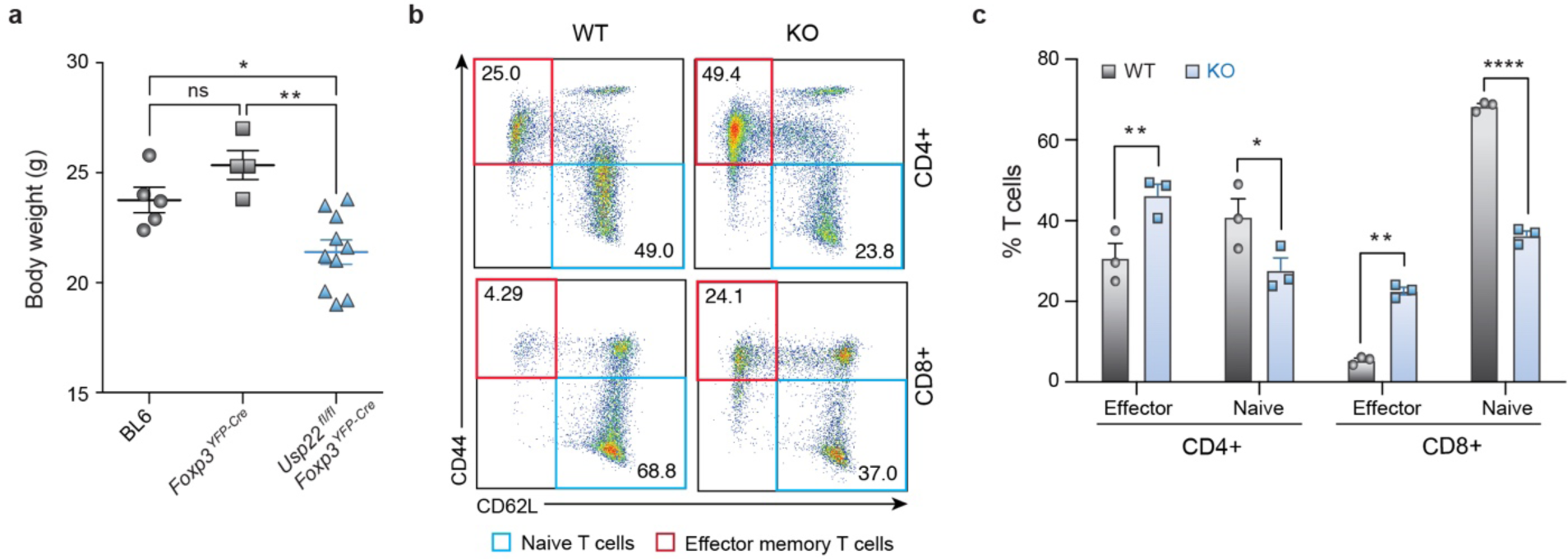
Autoimmune Inflammation in Treg-specific Usp22 Knockout Mice. a) Body weight differences between Usp22 WT and KO littermate mice. b) Representative flow cytometry analysis of CD44 and CD62L expression in splenic CD4^+^ and CD8^+^ T cells from 7-month-old WT and KO mice. Numbers in quadrants indicate percentage of each cell population. c) The frequency and absolute number of splenic effector memory-like T cells (CD44^hi^CD62L^lo^) and naïve T cells (CD44^lo^CD62L^hi^) of 7-month-old WT and KO mice were summarized. Data are mean ±SEM and are representative of three independent experiments (n=3). *P < 0.05, **P < 0.01, ****P < 0.0001.

**Extended Data Figure 7.**
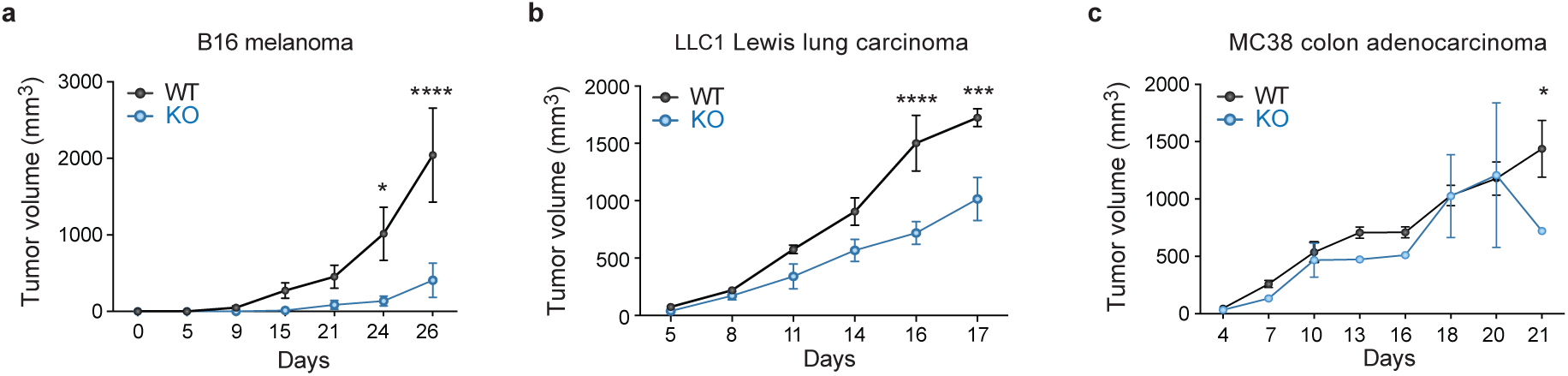
Tumor Growth is Inhibited in Treg-specific Usp22 Knockout Mice in Multiple Cancers. a) Mice were subcutaneously inoculated with 5^4^ B16 melanoma cells b) 10^6^ LLC1 Lewis lung carcinoma cells or c) 10^6^ MC38 colon adenocarcinoma cells. Tumor volumes were measured every 2-3 days by scaling along 3 orthogonal axes (x, y, and z) and calculated as (xyz)/2. *P < 0.05, **P < 0.01, ***P < 0.001, ****P < 0.0001.

## Methods

### Mice for Screen

B6 *Foxp3*^*GFP-Cre*^ mice^32^ were crossed with B6 *Rosa26*^*LSL-RFP*^ reporter mice^33^ as previously described^34^ to generate the Foxp3 fate reporter mice. These mice were then crossed to B6 constitutive Cas9-expressing mice^35^ to generate the *Foxp3*^*GFP-Cre*^*Rosa26*^*LSL-RFP*^*Cas9* mice used for the CRISPR screen. For the arrayed validation experiments, B6 *Foxp3*^*EGFP*^ knock-in mice^36^ that were obtained from Jackson Laboratories (Strain No. 006772) were used. These mice were maintained in the UCSF specific-pathogen-free animal facility in accordance with guidelines established by the Institutional Animal Care and Use Committee and Laboratory Animal Resource Center.

### Isolation and Culture of Primary Mouse Tregs for Screen and Validation

Spleens and peripheral lymph nodes were harvested from mice and dissociated in 1x PBS with 2% FBS and 1 mM EDTA. The mixture was then passed through a 70-µm filter. CD4^+^ T cells were isolated using the CD4^+^ Negative Selection Kit (StemCell Technologies, Cat# 19752) followed by fluorescence-activated cell sorting. For the prescreen sort, Tregs were gated on lymphocytes, live cells, CD4^+^, CD62L^+^, RFP^+^, Foxp3-GFP^+^ cells. For the arrayed validation experiments, Tregs were gated on lymphocytes, live cells, CD4^+^, Foxp3-GFP^+^ cells. Sorted Tregs were cultured in complete DMEM, 10% FBS, 1% pen/strep + 2000U hIL-2 in 24 well plates at 1 million cells/mL. Tregs were stimulated using CD3/CD28 Mouse T-Activator Dynabeads (Thermo Fisher, Cat# 11456D) at a ratio of 3:1 beads to cells for 48 hours. Cells were split and media was refreshed every 2-3 days.

### Pooled sgRNA Library Design and Construction

For the cloning of the targeted library, we followed the custom sgRNA library cloning protocol as previously described^37^. We utilized a MSCV-U6-sgRNA-IRES-Thy1.1 backbone (gifted from the Bluestone Lab). To optimize this plasmid for cloning the library, we first replaced the sgRNA with a 1.9kb stuffer derived from the lentiGuide-Puro plasmid (Addgene, plasmid# 52963) with flanking BsgI cut sites. This stuffer was excised using the BsgI restriction enzyme (NEB, Cat# R0559) and the linear backbone was gel purified (Qiagen, Cat# 28706). We designed a targeted library to include all genes matching Gene Ontology for “Nucleic Acid Binding Transcription Factors”, “Protein Binding Transcription Factors”, “Involved in Chromatin Organization” and “Involved in Epigenetic Regulation.” Genes were then selected based on those that have the highest expression levels across any mouse CD4 T cell subset as defined by Stubbington et al^38^. In total, we included 493 targets with 4 guides per gene, and 28 non-targeting controls. Guides were subsetted from the Brie sgRNA library^6^, and the pooled oligo library was ordered from Twist Bioscience to match the vector backbone. Oligos were PCR amplified and cloned into the modified MSCV backbone by Gibson assembly as described by Joung et al^37^. The library was amplified using Endura ElectroCompetent Cells following the manufacturer’s protocol (Endura, Cat# 60242-1). All oligos included in the library and primer sequences are listed in **<u>Supplementary Table 2</u>**.

### Retrovirus Production

Platinum-E (Plat-E) Retroviral Packaging cells (Cell Biolabs, Inc., Cat# RV-101) were seeded at 10 million cells in 15 cm poly-L-Lysine coated dishes 16 hours prior to transfection and cultured in complete DMEM, 10% FBS, 1% pen/strep, 1 µg/mL puromycin and 10 µg/mL blasticidin. Immediately before transfection, the media was replaced with antibiotic free complete DMEM, 10% FBS. The cells were transfected with the sgRNA transfer plasmids (MSCV-U6-sgRNA-IRES-Thy1.1) using TransIT-293 transfection reagent per the manufacturer’s protocol (Mirus, Cat# MIR 2700). The following morning, the media was replaced with complete DMEM, 10% FBS, 1% pen/strep. The viral supernatant was collected 48 hours post transfection and filtered through a 0.45 μm, polyethersulfone sterile syringe filter (Whatman, Cat# 6780-2504), to remove cell debris. The viral supernatant was aliquoted and stored until use at −80°C.

### Retroviral Transduction

Tregs were stimulated as described above for 48-60 hours. Cells were counted and seeded at 3 million cells in 1 mL of media with 2x hIL-2 into each well of a 6 well plate that was coated with 15 µg/mL of RetroNectin (Takara, Cat# T100A) for 3 hours at room temperature and subsequently washed with 1x PBS. Retrovirus was added at a 1:1 v/v ratio (1 mL) and plates were centrifuged for 1 hour at 2000g at 30°C and placed in the incubator at 37°C overnight. The next day, half (1 mL) of the 1:1 retrovirus to media mixture was removed from the plate and 1 mL of fresh retrovirus was added. Plates were immediately centrifuged for 1 hour at 2000g at 30°C. After the second spinfection, cells were pelleted, washed, and cultured in fresh media.

### Foxp3 Intracellular Stain and Post-Screen Cell Collection

Tregs were collected from their culture vessels 8 days after the second transduction and centrifuged for 5 min at 300g. Cells were first stained with a viability dye at a 1:1,000 dilution in 1× PBS for 20 min at 4°C, then washed with EasySep Buffer (1× PBS, 2% FBS, 1 mM EDTA). Cells were then resuspended in the appropriate surface staining antibody cocktail and incubated for 30 min at 4°C, then washed with EasySep Buffer. Cells were then fixed, permeabilized, and stained for transcription factors using the Foxp3 Transcription Factor Staining Buffer Set (eBioscience, cat. 00-5523-00) according to the manufacturer’s instructions. Antibody staining panels are listed in **<u>Supplementary Table 3</u>**. For the CRISPR screen, Foxp3^high^ and Foxp3^low^ populations were isolated using fluorescence-activated cell sorting by gating on lymphocytes, live cells, CD4^+^ and gating on the highest 40% of Foxp3-expressing cells (Foxp3^high^) and lowest 40% of Foxp3-expressing cells (Foxp3^low^) by endogenous Foxp3 intracellular staining. Over 2 million cells were collected for both sorted populations to maintain a library coverage of at least 1,000 cells per sgRNA (1000x).

### Isolation of Genomic DNA from Fixed Cells

After cell sorting and collection, genomic DNA (gDNA) was isolated using a protocol specific for fixed cells. Cell pellets were resuspended in cell lysis buffer (0.5% SDS, 50 mM Tris, pH 8, 10 mM EDTA) with 1:25 v/v of 5M NaCl to reverse crosslinking and incubated at 66°C overnight. RNase A (10 mg/mL) was added at 1:50 v/v and incubated at 37°C for 1 hour. Proteinase K (20 mg/mL) was added at 1:50 v/v and incubated at 45°C for 1 hour. Phenol:Chloroform:Isoamyl Alcohol (25:24:1) was added to the sample 1:1 v/v and transferred to a phase lock gel light tube (QuantaBio, Cat# 2302820), inverted vigorously and centrifuged at 20,000g for 5 mins. The aqueous phase was then transferred to a clean tube and NaAc at 1:10 v/v, 1 µl of GeneElute-LPA (Sigma, Cat# 56575), and isopropanol at 2.5:1 v/v were added. The sample was vortexed, and incubated at −80°C until frozen solid. Then thawed and centrifuged at 20,000g for 30 mins. The cell pellet was washed with 500 µl of 75% EtOH, gently inverted and centrifuged at 20,000g for 5 mins, aspirated, dried, and resuspended in 20 µl TE buffer.

### Preparation of Genomic DNA for Next Generation Sequencing

Amplification and bar-coding of sgRNAs for the cell surface sublibrary was performed as previously described^39^ with some modifications. Briefly, after gDNA isolation, sgRNAs were amplified and barcoded with TruSeq Single Indexes using a one-step PCR. TruSeq Adaptor Index 12 (CTTGTA) was used for the Foxp3^low^ population and TrueSeq Adaptor Index 14 (AGTTCC) was used for the Foxp3^high^ population. Each PCR reaction consisted of 50μL of NEBNext Ultra II Q5 Master Mix (NEB, Cat# M0544), 1μg of gDNA, 2.5μL each of the 10μM forward and reverse primers, and water to 100μL total. The PCR cycling conditions were: 3 minutes at 98°C, followed by 10 seconds at 98°C, 10 seconds at 62°C, 25 seconds at 72°C, for 26 cycles; and a final 2 minute extension at 72°C. After the PCR, the samples were purified using Agencourt AMPure XP SPRI beads (Beckman Coulter, cat #A63880) per the manufacturer’s protocol, quantified using the Qubit ssDNA high sensitivity assay kit (Thermo Fisher Scientific, cat #Q32854), and then analyzed on the 2100 Bioanalyzer Instrument. Samples were then sequenced on an Illumina MiniSeq using a custom sequencing primer. Primer sequences are listed in **<u>Supplementary Table 2</u>**.

### Pooled CRISPR Screen Pipeline

Primary Tregs were isolated from the spleen and lymph nodes of three male Foxp3-GFP-Cre/Rosa26-RFP/Cas9 mice aged 5-7 months old, pooled together, and stimulated for 60 hours. Cells were then retrovirally transduced with the sgRNA library and cultured at a density of 1 million cells/ml continually maintaining a library coverage of at least 1,000 cells per sgRNA. Eight days after the second transduction, cells were sorted based on Foxp3 expression defined by intracellular staining. Genomic DNA was harvested from each population and the sgRNA-encoding regions were then amplified by PCR and sequenced on an Illumina MiniSeq using custom sequencing primers. From this data, we quantified the frequencies of cells expressing different sgRNAs in each in each population (Foxp3^high^ and Foxp3^low^) and quantified the phenotype of the sgRNAs, which we have defined as Foxp3 stabilizing (enriched in Foxp3^high^) or Foxp3 destabilizing (enriched in Foxp3^low^).

### Analysis of Pooled CRISPR Screen

Analysis was performed as previously described^40^. To identify hits from the screen, we used the MAGeCK software to quantify and test for guide enrichment^7^. Abundance of guides was first determined by using the MAGeCK “count” module for the raw fastq files. For the targeted libraries, the constant 5’ trim was automatically detected by MAGeCK. To test for robust guide and gene-level enrichment, the MAGeCK “test” module was used with default parameters. This step included median ratio normalization to account for varying read depths. We used the non-targeting control guides to estimate the size factor for normalization, as well as to build the mean-variance model for null distribution, which was used to find significant guide enrichment. MAGeCK produced guide-level enrichment scores for each direction (i.e. positive and negative) which were then used for alpha-robust rank aggregation (RRA) to obtain gene-level scores. The p-value for each gene was determined by a permutation test, randomizing guide assignments and adjusted for false discovery rates by the Benjamini–Hochberg method. Log2 fold change (LFC) was also calculated for each gene, defined throughout as the median LFC for all guides per gene target. Where indicated, LFC was normalized to have a mean of 0 and standard deviation of 1 to obtain the LFC Z-score. Output files from the MAGeCK analysis are in **<u>Supplementary Table 1</u>**.

### Arrayed Cas9 Ribonucleotide Protein (RNP) Preparation and Electroporation

RNPs were produced by complexing a two-component gRNA to Cas9, as previously described^17^. In brief, crRNAs and tracrRNAs were chemically synthesized (IDT), and recombinant Cas9-NLS were produced and purified (QB3 Macrolab). Lyophilized RNA was resuspended in Nuclease-free Duplex Buffer (IDT, Cat# 1072570) at a concentration of 160 µM, and stored in aliquots at − 80 °C. crRNA and tracrRNA aliquots were thawed, mixed 1:1 by volume, and annealed by incubation at 37 °C for 30 min to form an 80 µM gRNA solution. Recombinant Cas9 was stored at 40 µM in 20 mM HEPES-KOH, pH 7.5, 150 mM KCl, 10% glycerol, 1 mM DTT, were then mixed 1:1 by volume with the 80 µM gRNA (2:1 gRNA to Cas9 molar ratio) at 37 °C for 15 min to form an RNP at 20 µM. RNPs were electroporated immediately after complexing. RNPs were electroporated 3 days after initial stimulation. Tregs were collected from their culture vessels and centrifuged for 5 min at 300g, aspirated, and resuspended in the Lonza electroporation buffer P3 using 20 µl buffer per 200,000 cells. 200,000 Tregs were electroporated per well using a Lonza 4D 96-well electroporation system with pulse code EO148 (mouse) or EH115 (human). Immediately after electroporation, 80 μL of pre-warmed media was added to each well and the cells were incubated at 37°C for 15 minutes. The cells were then transferred to a round-bottom 96-well tissue culture plate and cultured in complete DMEM, 10% FBS, 1% pen/strep + 2000U hIL-2 at 200,000 cells/well in 200 µl of media.

### Isolation and Culture of Human Treg Cells

Primary human Treg cells for all experiments were sourced from residuals from leukoreduction chambers after Trima Apheresis (Blood Centers of the Pacific) under a protocol approved by the UCSF Committee on Human Research (CHR# 13-11950). Peripheral blood mononuclear cells (PBMCs) were isolated from samples by Lymphoprep centrifugation (StemCell, Cat #07861) using SepMate tubes (StemCell, Cat# 85460). CD4^+^ T cells were isolated from PBMCs by magnetic negative selection using the EasySep Human CD4^+^ T Cell Isolation Kit (StemCell, Cat# 17952) and Tregs were then isolated using fluorescence-activated cell sorting by gating on CD4^+^, CD25^+^, CD127^low^ cells. After isolation, cells were stimulated with ImmunoCult Human CD3/CD28/CD2 T Cell Activator (StemCell, Cat# 10970) per the manufacturer’s protocol and expanded for 9 days. Cells were cultured in complete RPMI media, 10% FBS, 50mM 2-mercaptoethanol and 1% pen/strep with hIL-2 at 300U/mL at 1 million cells/mL. After expansion, Tregs were restimulated in the same way for 48h before RNP electroporation.

### Generation of Treg-Specific Usp22 Knockout Mice

Usp22 floxed mice were generated and used as recently reported^41^. The Usp22 target mouse embryonic stem cells from C57/B6 mice were purchased from the Wellcome Trust Sanger Institute. Blastocyst injections resulted in several chimeric mice with the capacity for germline transmission. Breeding of heterozygous mice yielded *Usp22*^*wt/wt*^, *Usp22*^*wt/targeted*^ but not *Usp22*^*targeted/targeted*^ mice due to the obligation of Usp22 expression by the neomycin selection and β-gal reporter cassette, which causes embryonic lethality^42^. We then bred *Usp22*^*wt/targeted*^ mice with Flp recombinase transgenic mice to delete the selection cassette, leading to the generation of *Usp22*^*wt/fl*^ mice, further breeding of which produced *Usp22*^*wt/wt*^, *Usp22*^*wt/fl*^ and *Usp22*^*fl/fl*^ mice without phenotypic abnormalities in expected Mendelian ratios (**Extended Data Figs. 3a, 3b**). Treg-specific Usp22-null mice were generated by breeding *Usp22*^*fl/fl*^ mice with *Foxp3*^*YFP-Cre*^ mice^18^. Additionally, C57BL/6 Rag-/-mice, SJL CD45.1 congenic mice were purchased from Jackson Laboratories. These mice were maintained and used at the Northwestern University mouse facility under pathogen-free conditions according to institutional guidelines and using animal study proposals approved by the institutional animal care and use committees. Unless stated otherwise, all figures are representative of experiments with healthy 6- to 8-week-old mice.

### Cell lines, Plasmids, Antibodies, and Reagents

Human embryonic kidney 293 cells (HEK293) were stored in the Fang lab and were cultured in DMEM containing 10% FBS. EG7 lymphoma, MC38 colon cancer, LLC1-OVA lung carcinoma and B16-SIY melanoma cell lines were provided by Dr. Bin Zhang and used for tumor models as previously reported^43^. Myc-Usp22, Myc-Usp22(C185A), FLAG-Foxp3 and HA-ubiquitin expression plasmids and their tagged vectors were constructed and stored in the Fang lab. Antibodies used for Western blots, Co-IPs and flow cytometry are listed in **<u>Supplementary Table 3</u>**. PMA (phorbol 12-myristate 13-acetate), ionomycin, and Cycloheximide (CHX) were purchased from Sigma. Monesin was from eBioscience.

### Cell Isolation and Flow Cytometry for Analysis of Usp22 KO Mice

Peripheral T cells were isolated from mouse spleen by a CD4^+^ T-cell negative (Stem Cell) or positive selection kit (Invitrogen). Enriched CD4^+^ T cells were further sorted for either YFP^+^ (Foxp3^+^) T cells or CD25^-^CD44^-lo^CD62L^hi^ naïve T cells by FACSAria (BD Bioscience). Purity of sorted cells was > 99%. Lymphocytes isolated from the intestinal lamina propria were acquired by following previously described methods^44^. To isolate tumor-infiltrating lymphocytes, tumors were cut into small fragments and digested by collagenase D (Sigma) and DNase (Sigma) for 1h at room temperature.

Flow cytometry was done with a FACSCanto II. Samples were initially incubated with anti-CD16/32 antibodies to block antibody binding to Fc receptor. Single-cell suspensions were stained with relevant antibodies according to each protocol and then washed twice with cold PBS containing 3% FBS. For intracellular cytokine staining, cells were first stimulated for 4-5 h with 20 ng/ml PMA plus 0.5μM ionomycin in the presence of monesin (10 μg/ml), and the fixed cells were incubated with the appropriate antibodies and analyzed by flow cytometry. Data were analyzed with FlowJo software.

### Tumor Models

Cultured cancer cells were trypsinized and washed once with PBS. Tumor cells (1×10^6^ EG7 cells, LLC1 cells or MC38 cells or 5×10^4^ B16 melanoma cells) in suspension were subcutaneously injected into WT or *Usp22*^*fl/fl*^*Foxp3*^*YFP-Cre*^ eight to ten week-old mice. Tumors were measured every 2-3 days by measuring along 3 orthogonal axes (x, y, and z) and calculated as (xyz)/2 as recently reported^43^.

### *In Vitro* Treg Suppression Assay

Naïve CD4^+^CD25^-^ T cells (5×10^4^) labeled with eFluor 670 cell proliferation dye were used as responder T cells and cultured in 96-well U-bottom plate for 72h together with increasing ratio of sorted YFP+ Treg cells from WT or *Usp22*^*fl/fl*^ *Foxp3*^*YFP-Cre*^ mice in the presence of irradiated splenocytes depleted of T cells (5×10^4^) plus anti-CD3 (2 μg/ml). The suppressive function of Treg cells was determined by measurement of the proliferation of activated CD4^+^ and CD8^+^ effector T cells on the basis of eFluor 670 cell proliferation dye dilution as reported^45^.

### Induced Treg Differentiation

We isolated 0.5×10^6^ CD4^+^CD25^-^ naïve T cells from splenocytes of WT or *Usp22*^*fl/fl*^*Foxp3*^*YFP-Cre*^ mice and cultured them in 24-well plates coated with 3 μg/ml anti-CD3 and 5μg/ml anti-CD28 antibodies for 5 days. For iTreg cell polarization, the cultures were supplemented with IL-2 (5 ng/ml), anti-IFN-γ (2 μg/ml), anti-IL-4 (2 μg/ml) and TGF-β (at 2, 5 or 10 ng/ml). Cytokines were purchased from Peprotech.

### Quantitative PCR (qPCR)

RNA was extracted from sorted YFP^+^ Tregs and qPCR was performed following the manufacturer’s protocol using gene-specific primer sets (**<u>Supplementary Table 2</u>**).

### Histology

Mouse tissues were fixed in 10% formalin and embedded in paraffin. 4μm sections were stained with hematoxylin and eosin.

### Co-Immunoprecipitation and Western blot

Co-IPs and Western blots were performed as previously described^46^. Cells were collected and resuspended in RIPA buffer (Millipore, Cat# 20-188) with protease inhibitors (Roche, Cat# 36363600) and incubated on ice for 30 min. Cells were centrifuged (12000g for 10 min) at 4 °C and the cell debris was discarded. The supernatant was incubated with protein G-sepharose beads at 4 °C for 30 min and then with the indicated antibody (1 μg/test) for 2 h followed by incubation with protein G-sepharose beads overnight with rotation at 4 °C. The cells were then washed 5 times with RIPA buffer and the protein G-sepharose beads were dissolved with loading buffer and boiled for 5 min. Supernatants were subjected to SDS-PAGE gel and transferred to nitrocellulose membrane. With blocking with 5% (w/v) skim milk in TBS-T buffer, the membrane was incubated overnight with indicated primary antibodies at 4°C followed by HRP-conjugated secondary antibody or with HRP conjugated primary antibodies (**<u>Supplementary Table 3</u>**). Membranes were then developed with enhanced chemiluminescence (ECL).

### Ubiquitination Assay

Flag-Foxp3 and HA-ubiquitin plasmids were co-transfected into HEK293 cells using Turbofect Transfection Reagent (Cat# R0532) along with either Myc-empty vector, Myc-Usp22, or the catalytically inactive mutant Myc-Usp22C185A (C>A), where the conserved cysteine (C) residue in the C19 peptidase domain was replaced by an alanine (A) residue. After 48 hours, cells were collected, immunoprecipitated with anti-Flag to pull down Foxp3, and immunoblotted for HA-ubiquitin to assess Foxp3 ubiquitination in the presence or absence of functional Usp22. Whole cell lysate (WCL) controls were immunoblotted with HRP-conjugated Myc and HRP-conjugated Flag to show transfection efficiency.

### Chromatin Immunoprecipitation (ChIP)

T cells were polarized using Treg polarizing conditions in a 24-well plate for 3 days, and 3 million cells were used per immunoprecipitation. Cells were fixed in 37% formaldehyde for 10 min at 37°C. Glycine was added to a final concentration of 0.125 M, and the incubation was continued for an additional 5 min at room temperature. Cells were washed twice with ice-cold phosphate-buffered saline with 1x Protease Inhibitor cocktail (Roche, Cat# 36363600). Millipore ChIP Assay Kit (LOT 3154126) was used for the remainder of the protocol. Cells were resuspended in 1 ml of SDS lysis buffer (Millipore, Cat# 20-163) with protease inhibitors and set on ice for 15 minutes. Samples were then sonicated at the medium setting (308/608) for 7 minutes. Samples were centrifuged at 14 000 rpm at 4°C for 10 min. After removal of an input control (whole-cell extract), supernatants were diluted 10-fold in ChIP dilution buffer (Millipore, Cat# 20-153), and 1x protease inhibitor. 40 uL of Salmon Sperm DNA/Protein A Agarose-50% (Millipore, Cat #16-157C) for 30 min spinning at 4°C. Agarose pelleted out with brief centrifugation and supernatant moved to a new tube. Samples were incubated with either 4 ul of antibody rabbit anti-IgG (Cell Signaling, Cat# 2729), rabbit anti-USP22 (Abcam, ab195289), ub-Histone H2A (Cell Signaling, Cat# 8204S), and ub-Histone H2B (Cell Signaling, Cat# 5546P) overnight at 4°C rotation. Samples then incubated with 30 ul of of Salmon Sperm DNA/Protein A Agarose-50% for 1 hour at 4°C with rotation. Agarose was pelleted and placed at rotation for five minutes with Low Salt Immune Complex Wash Buffer (Millipore, Cat #20-154), then pelleted. High Salt Immune Complex Wash Buffer (Millipore, Cat #20-155) was added to the pellet and the sample was spun for five minutes at 4°C then pelleted. The agarose was re-suspended in LiCl Immune Complex Wash Buffer and placed at rotation for five minutes at 4°C. The samples were then spun down and re-suspended in 1X TE (Millipore, Cat #20-157) and placed at rotation at room temperature for 5 minutes (repeated once more). The sample was then pelleted and resuspended in 100uL of elution buffer (1%SDS, 0.1M NaHCO3 in water) and placed at rotation for 10 minutes at room temperature. Sample was spun down and supernatant was saved, and step was repeated. 10uL of 5M NaCl was added to the combined eluates and to the imput starting material at heated at 65°C overnight. 0.5 M EDTA, 1M Tris-HCl (pH 6.5) and 10mg/mL of protein kinase were added to the samples and incubated at 45°C for one hour. DNA was recovered using a PCR purification kit (Qiagen, Cat #28004).

### RNA sequencing

YFP+ Treg cells were sorted from spleen and LN of WT or *Usp22*^*fl/fl*^*Foxp3*^*YFP-Cre*^ mice (n=5 per group) and total RNA was isolated from 1×10^6^ cells per sample using an RNeasy Mini Kit (Qiagen, Cat# 74104) as previously described^47^. ERCC ExFold RNA Spike-In Mixes (Thermo Fisher, Cat# 4456739) were then added to total RNA for each sample. WT samples were spiked with Mix #1, while KO samples were spiked with Mix# 2; all samples were spiked with 8.16ul of 1:1000 diluted spike-in mix. Total RNA was then provided to the Functional Genomics Laboratory at UC Berkeley where RNA-seq libraries were prepared by Oligo dT enrichment followed by a stranded Illumina library prep protocol with the KAPA mRNA HyperPrep kit (Kapa Biosystems, KK8580). Libraries were checked for quality on an AATI Fragment analyzer (Agilent, DNF-935-1000), quantified using the Illumina Quant Universal qPCR Mix (Kapa Biosystems, KK4824), and pooled evenly at 3nM. The Vincent J. Coates Genomics Sequencing Laboratory at UC Berkeley then performed one lane of 150bp paired-end Illumina, HiSeq4000 sequencing followed by demultiplexing and bclfile to fastq conversion using Illumina bcl2fastq v2.19 software (Illumina). Reads were mapped to the GRCm38.p6 assembly (Ensembl annotation) using kallisto v0.45 with default parameters^48^. Transcript-level abundance estimates were collapsed to create gene-level count matrices and normalized to the ERCC spike-ins using loess regression^49^. Differentially expressed genes were then detected using DESeq2 with default parameters^50^. Data from this experiment is included in **<u>Supplementary Table 4</u>**.

### Adoptive transfer model of colitis

Naïve T cells (CD4^+^CD25^-^CD44^lo^CD62L^hi^) were sorted from congenic CD45.1 B6.SJL mice and YFP+ Treg cells were sorted from WT or *Usp22*^*fl/fl*^*Foxp3*^*YFP-Cre*^ mice. Rag1^-/-^ mice were given intraperitoneal injection of naïve T cells (4×10^5^) alone or in combination with WT or Usp22 KO Treg cells (2×10^5^). After T cell reconstitution, mice were weighed weekly and monitored for clinical of signs of disease. Mice were sacrificed when their body weight decreased 20%. At cessation, colons were harvested for measurement and histology and flow cytometry.

### Induced experimental autoimmune encephalomyelitis (EAE)

Eight to ten week-old WT or *Usp22*^*fl/fl*^*Foxp3*^*YFP-Cre*^ mice were subcutaneously injected with 200µg of MOG_33-55_ peptide (Genemed Synthesis). The MOG_33-55_ peptide was emulsified in complete Feund’s adjuvant (CFA) which contained 200µg of *Mycobacterium tuberculosis* H37Ra (Difco). The mice were then subsequently intraperitoneally injected with 200 ng of pertussis toxin (List Biological Laboratories) on day 0 and day 2. Clinical signs of EAE were assessed daily. Scores were given as follows: 0, no sign of disease; 2, limp tail, 3, hind leg weakness or limp; 3, partial back limb paralysis; 4, complete hind limb paralysis; 5, total limb paralysis.

### Data availability

Data from the screen (Fig. 1) and RNA sequencing (Fig. 3) are provided here in Supplementary Tables 1 and 4.

### Code availability

All custom code used in this manuscript will be made available before publication or on request.

---

**<u>Supplementary Table 1</u>**: Screen data. This file includes result files from MAGeCK analysis for sgRNA and gene level enrichment and normalized and raw count files.

**<u>Supplementary Table 2</u>**: Synthetic oligos used in this study - sgRNA library, primers and crRNA for RNP arrays.

**<u>Supplementary Table 3</u>**: Antibodies used in this study.

**<u>Supplementary Table 4</u>**: Differentially expressed genes and raw counts from RNA sequencing.

## Acknowledgements

We thank all members of the Marson lab as well as Mark S. Anderson, K. Mark Ansel, Chun J. Ye, Kathrin Schumann and Luke Gilbert for helpful suggestions and technical advice. We thank Jacob Freimer, Siddharth Raju and Eric Guo for helpful advice and assistance with the RNA-seq analysis pipeline. We thank Vinh Nguyen, Victoria Tobin, Ryan Apathy, Jonathan Woo, Michelle Nguyen and UCSF Flow Cytometry Core for technical assistance. We thank Sarah Pyle for assistance with graphics. We thank David Nguyen for critical reading of the manuscript. J.T.C. is supported by the National Science Foundation Graduate Research Fellowship Program grant 1650113. The Marson lab has received gifts from J. Aronov, G. Hoskin, K. Jordan, B. Bakar, the Caufield family and has received funds from the Innovative Genomics Institute (IGI) and the Parker Institute for Cancer Immunotherapy (PICI). A.M. holds a Career Award for Medical Scientists from the Burroughs Wellcome Fund and is an investigator at the Chan Zuckerberg Biohub. This work used the Vincent J. Coates Genomics Sequencing Laboratory at UC Berkeley, supported by NIH S10 OD018174 Instrumentation Grant and the UCSF Flow Cytometry Core, supported by the Diabetes Research Center grant NIH P30 DK063720.

## Author Contributions

Conceptualization: J.T.C., E.M., E.S., Yu.Z., Z.S., F.V.G., J.A.B., A.M., and D.F. Methodology: J.T.C., E.S., T.L.R. Investigation: J.T.C., E.M., E.S., Yu.Z., O.S., Y.X., D.R.S., Ya.Z., T.L.R, S.C., Z.L., I.A.V., G.Y.P., Y.L., and I.I. Resources: J.A.B., A.M., and D.F. Formal analysis: E.S., J.T.C., Software: E.S., Data Curation: J.T.C., E.S., Supervision: B.Z., Y.L., F.V.G., J.A.B., A.M., and D.F. Funding acquisition: J.T.C., E.M., A.M., and D.F. Writing – original draft preparation: J.T.C., E.M., Yu.Z. A.M., and D.F. Writing – review and editing: J.T.C., E.M., Z.S., F.V.G., J.A.B., A.M., and D.F.

